# Single Molecule Studies and Kinase Activity Measurements Reveal Regulatory Interactions between the Master Kinases Phosphoinositide-Dependent-Kinase-1 (PDK1), Protein Kinase B (AKT1/PKB) and Protein Kinase C (PKCα)

**DOI:** 10.1101/2021.06.25.449974

**Authors:** Moshe T. Gordon, Brian P. Ziemba, Joseph J. Falke

**Author notes:** Corresponding Author: Joseph J. Falke.

## Abstract

Leukocyte migration is controlled by a leading edge chemosensory pathway that generates the regulatory lipid PIP3, a growth signal, thereby driving leading edge expansion up attractant gradients toward sites of infection, inflammation, or tissue damage. PIP_3_ also serves as an important growth signal in growing cells and oncogenesis. The kinases PDK1, AKT1/PKB and PKCα are key components of a plasma membrane-based PIP_3_ and Ca^2+^ signaling circuit that regulates these processes. PDK1 and AKT1 are recruited to the membrane by PIP_3_, while PKCα is recruited to the membrane by Ca^2+^. All three of these master kinases phosphoregulate an array of protein targets. For example, PDK1 activates AKT1, PKCα and other AGC kinases by phosphorylation at key sites. PDK1 is known to form PDK1:AKT1 and PDK1:PKCα heterodimers stabilized by a PIF interaction between the PDK1 PIF pocket and the PIF motif of the AGC binding partner. Here we present the first, to our knowledge, single molecule studies of full length PDK1 and AKT1 on target membrane surfaces, as well as their interaction with full length PKCα. The findings show that membrane-bound PDK1:AKT1 and PDK1:PKCα heterodimers form under physiological conditions, and are stabilized by PIF interaction. PKCα exhibits 8-fold higher PDK1 affinity than AKT1, thus PKCα competitively displaces AKT1 from PDK1:AKT1 heterodimers. Ensemble activity measurements under matched conditions reveal that PDK1 activates AKT1 via a *cis* mechanism by phosphorylating an AKT1 molecule in the same PDK1:AKT1 heterodimer, while PKCα acts as a competitive inhibitor of this phosphoactivation reaction by displacing AKT1 from PDK1. Overall, the findings provide new insights into molecular and regulatory interactions of the three master kinases on their target membrane, and suggest that the recently described tumor suppressor activity of PKC may arise from its ability to downregulate PDK1-AKT1 phosphoactivation in the PIP3-PDK1-AKT1-mTOR pathway linked to cell growth and oncogenesis.

**STATEMENT OF SIGNIFICANCE:** This work investigates three master kinases that play central roles in guiding white blood cell migration to sites of infection, inflammation or tissue damage. More broadly, the same kinases help regulate production of a cell growth signal, and may trigger cancer when dysregulated. Using powerful single molecule methods, the work detects and analyzes the interactions between the three purified kinases on their target membrane surface. The findings reveal functionally important differences between pairwise binding affinities of different binding partners. Additional studies reveal that the highest affinity kinase can disrupt and inhibit the activated complex formed by association of the other two kinases. Such inhibition is proposed to help prevent cancer by limiting growth signal production by the activated complex.

## INTRODUCTION

The chemosensory pathway of macrophages and other leukocytes is located on the leading edge pseudopod membrane that controls the direction of cell migration by expanding up attractant concentration gradients. In the regulatory core of this pathway, signals from attractant-activated receptors, G proteins and a local Ca^2+^ influx activate multiple isoforms of class I PI3K, a lipid kinase that phosphorylates the substrate lipid PIP_2_ to generate the essential signaling lipid PIP_3_(1–13). **Figure 1** outlines the core pathway in which several key activation mechanisms are now understood at the molecular level. Receptor tyrosine kinases generate phosphotyrosine (PiTyr) residues on membrane-bound or soluble activator proteins (14–20), and these PiTyr residues activate PI3K by binding and displacing the inhibitory SH2 domains that block the membrane docking surface of the lipid kinase domain (18, 21). Other receptors, including GPCRs, generate signals that activate members of the Ras superfamily, which are small, membrane-anchored G protein that can stimulate PI3K by enhancing its membrane recruitment (4, 21–23). The local Ca^2+^ signal recruits protein kinase C (PKCα) to the leading-edge membrane where it phosphorylates the MARCKS protein, thereby displacing MARCKS from the PIP_2_ lipid it sequesters (11, 24–26). The resulting free PIP_2_ recruits additional PI3K to the membrane and also serves as the substrate for lipid kinase activity and production of the PIP_3_ signal. The signaling lipid PIP_3_ is a ubiquitous growth signal that plays an essential role in driving leukocyte pseudopod growth up an attractant gradient (reviewed in (24, 26)). The PIP_3_ signal recruits downstream PH domain proteins to the leading edge membrane, including 3-phospho-inositide-dependent protein kinase 1 (PDK1) (27) and RAC-alpha serine/threonine-protein kinase (AKT1, also known as protein kinase B or PKB) (28, 29). Both PDK1 and AKT1 undergo phosphoactivation on the membrane surface and stimulate multiple, essential downstream pathways (29–33). More broadly, in all growing cells PIP_3_ is a growth signal, and mutations that generate excessive PIP_3_ levels are often linked to cancer and other diseases. Thus Ras, PI3K and AKT1 are well established oncogenes, and a deeper molecular understanding of the core pathway is relevant not only to macrophage chemotaxis and innate immunity but is also central to cell growth and cancer biology (14, 16, 34–36).

**Figure 1.**
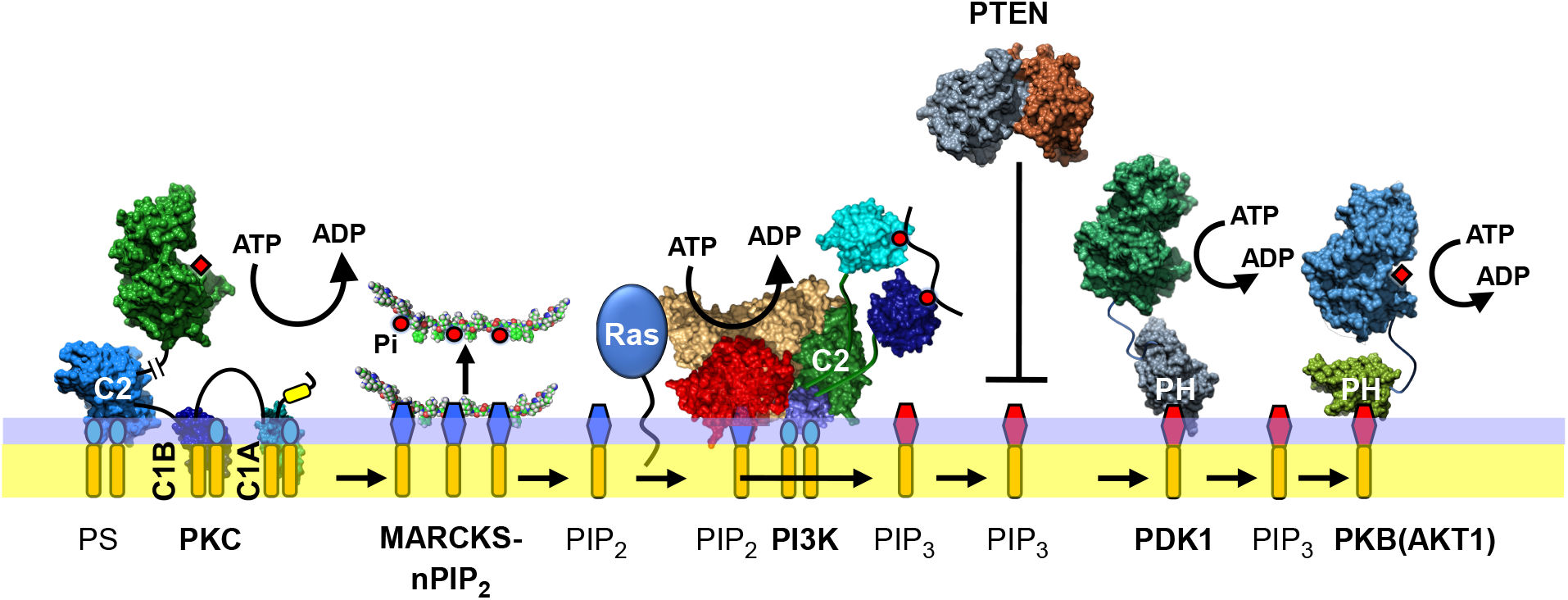
Regulatory core of the Ca^2+^/PIP_3_ signaling circuit on the leading edge membrane of chemotaxing leukocytes (reviewed in (25, 23)). Protein kinase C isoforms (including PKCα) integrate activating signals from Ca^2+^ and diacylglycerol then phosphorylate MARCKS, which releases sequestered PIP_2_ lipid. The resulting free PIP_2_ is the binding target and substrate for Class I isoforms of the lipid kinase phosphatidylinositol-3-kinase (PI3K). PI3K is activated by phosphotyrosines, often generated by receptor tyrosine kinases, and then is recruited to the plasma membrane by PIP_2_ and by a GTP-Ras superfamily isoform (if present and anchored to the membrane by one or more lipidations). Once activated and localized to the membrane, PI3K phosphorylates PIP_2_, generating the signaling lipid and growth signal PIP_3_. The resulting PIP_3_ recruits PDK1 and AKT1 to the leading-edge membrane and triggers their phosphoactivation reactions. In this study we quantify the affinity of PDK1:PKC and PDK1:AKT1 heterodimers and measure the effects of heterodimer formation on protein kinase activity.

PDK1 is a master protein kinase that phosphoactivates a wide array of crucial AGC kinases, including AKT1 and PKC isoforms such as PKCα (37, 38). Previous studies have described a “PIF pocket” cleft on PDK1 that binds a conserved PDK1-interacting fragment (PIF) motif on AKT1, PKCα and other AGC kinases (**Figure 2**). This PIF interaction is known to be essential for phosphoactivation (37, 39). Occupation of this cleft either by a protein, peptide, or small molecule causes an allosteric conformational change, allowing for full kinase activity of PDK1 (40).

**Figure 2.**
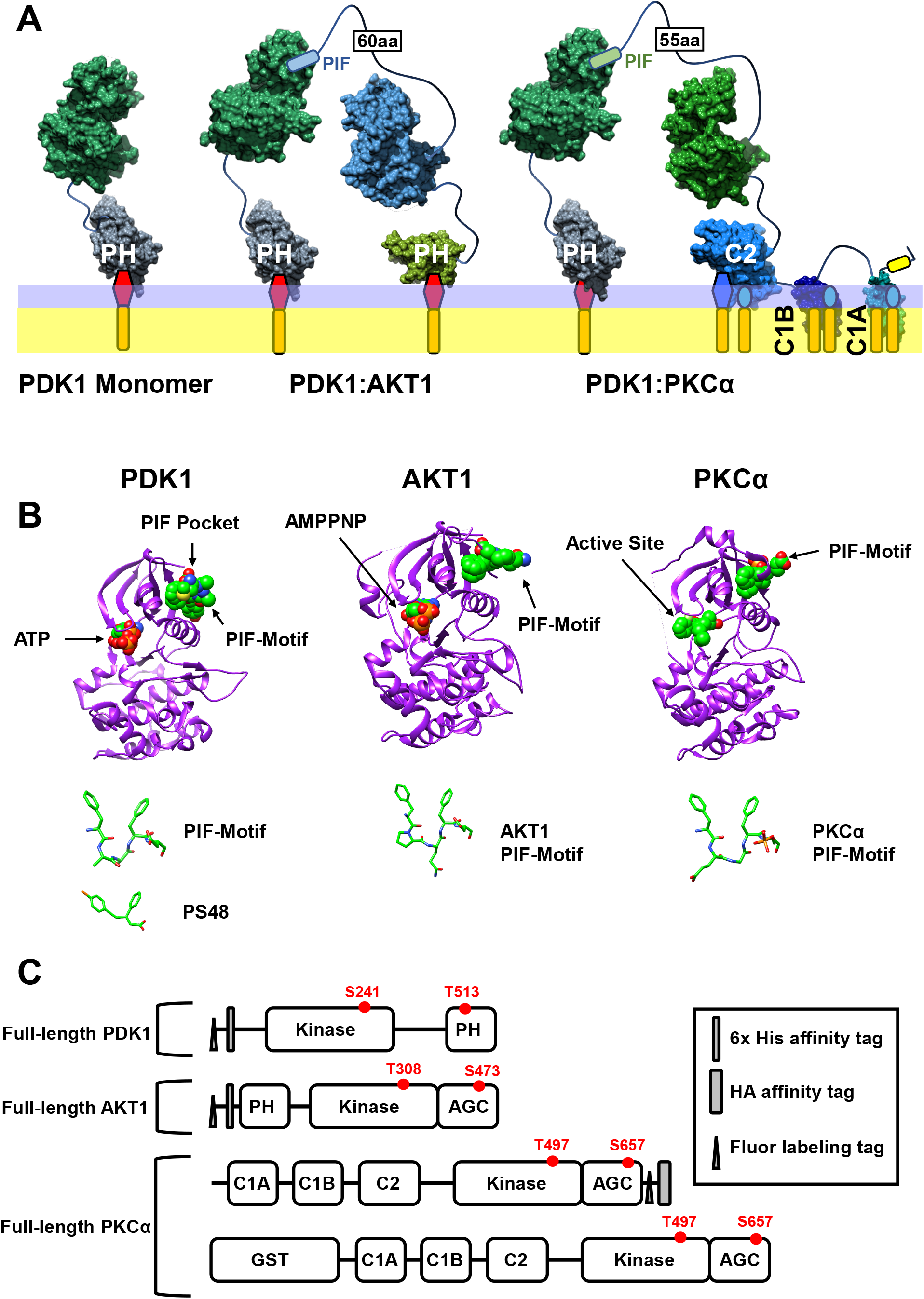
Schematic models and structures of the three master protein kinases studied herein, illustrating the PIF interactions that stabilize the detected heterodimers. **(A)** Schematic membrane-bound structures of the PDK1 monomer and the PIF-stabilized PDK1:AKT1 and PDK1:PKCα heterodimers. Membrane targeting is provided by a PIP_3_-specific PH domain (PDK1, AKT1) or by a PS-specific C2 domain (PKCα). All three possess a tethered kinase domain. The AGC kinases (AKT1, PKCα) also possess a largely unstructured AGC domain containing a PIF motif that binds to the PIF pocket on PDK1, thereby stabilizing heterodimer formation. This working model proposes that the AGC domain acts as a long, flexible tether within the PIF-stabilized heterodimer, enabling the PDK1 active site to access the distal activation loop of the substrate kinase (62). (**B**) Crystal structures of the three protein kinase domains: PDK1 occupied by ATP and a PIF peptide (left, PDB 4RRV); AKT1 occupied by AMPPNP and displaying its C-terminal PIF motif (middle, 4EKK); and PKCα occupied by inhibitor compound 28 (((R)-6-((3S,4S)-1,3-Dimethyl-piperidin-4-yl)-7-(2-fluoro-phenyl)-4-methyl-2,10-dihydro-9-oxa-1,2,4a-triaza-phenanthren-3-one)) and displaying its C-terminal PIF motif (right, 4RA4). Shown also are structural similarities between the small molecule PIF pocket inhibitor PS48 and representative PIF motifs. **(C)** Domain structures of the kinases showing relevant phosphorylation sites. Shown also are the ybbr fluor-labeling tag and the relevant purification tag for each construct. Two PKC constructs with equivalent activity were employed (see Methods).

Important mechanistic and functional questions remain unanswered regarding the regulatory interactions between PDK1 and its target AGC kinases on membrane surfaces. Many previous studies of the PIF interaction and PDK1 phosphoactivation of AGC kinases have used peptide substrates in solution, which have provided valuable insights, but which may not fully describe how the full length protein kinases are regulated on a cell membrane (32, 33, 37). PDK1 phosphoactivates AGC kinases by phosphorylating the activation loop adjacent to the AGC kinase active site cleft (27, 39). Previous studies which measured protein kinase activity of PDK1 on peptides used two classes of peptide substrates: peptides that contain both the activation loop phosphorylation site and the PIF motif on the same peptide (PIFtide), and peptides containing only the phosphorylation site with no PIF motif (t308tide). Much higher PDK1 protein kinase activity is observed when the substrate peptide contains both the activation loop phosphorylation site and a PIF motif (37). These data are consistent with a *cis* activation model for PDK1 activation of AGC kinases in which PDK1 phosphorylates an AGC kinase within the same PDK1:AGC heterodimer stabilized by the PIF interaction. On the other hand, as shown in **Figure 2,** the PIF interaction occurs on the surfaces of PDK1 and the AGC kinase distal to their active site clefts. Thus, a *cis* model requires considerable flexibility of the fulllength kinases in the activation reaction. Such flexibility would most likely originate from the substrate AGC kinase: both AKT1 and PKCα possesses a long (55+ residue), largely unstructured C-terminal AGC domain containing the PIF motif, as well as a flexible activation loop containing the target phosphorylation site. If sufficient flexibility is lacking, the distal geometry raises the possibility that PDK1 activation of AGC kinases could employ a *trans* mechanism in which activated PDK1 within the PDK1-AGC heterodimer phosphorylates a separate AGC monomer that collides with the heterodimer. The *cis* and *trans* models make contrasting, testable predictions about competition between AGC kinases. The *cis* model predicts that phosphactivation of one AGC kinase by PDK1 will be competitively inhibited by the binding of a different AGC kinase to PDK1, while the *trans* model predicts that such binding need not be inhibitory and could even be activating if a larger fraction of the PDK1 population is converted to PIF-activated heterodimers.

It follows that, in principle, competitive binding of AGC kinases to PDK1 could play an important regulatory role in the PI3K/PIP_3_ pathway, since PDK1 and its target AKT1 and PKCα enzymes are recruited to the same region of membrane by PIP_3_ and Ca^2+^ signals, respectively. As increasing PIP_3_ and Ca^2+^ levels raise the local densities of these membrane bound proteins, competition between AKT1 and PKCα for binding to PDK1 could in principle modulate the rate of phosphoactivation of the less prevalent AGC kinase.

In this study we employ single molecule methods to detect and quantify kinase-kinase interactions on a target membrane surface *in vitro*. The findings directly demonstrate that PDK1 forms stable, membrane-bound PDK1:AKT1 and PDK1:PKCα heterodimers via the PIF interaction. Titrations enable measurement of both heterodimer affinities and show that the heterodimers are formed under physiologically relevant conditions at nanomolar protein concentrations, with the PDK1:PKCα heterodimer exhibiting ~10-fold higher affinity than the PDK1:AKT1 heterodimer. The single molecule studies further reveal that the heterodimers are disrupted by a drug that binds to the PIF pocket (PS48), and that AKT1 and PKCα compete for binding to the PIF pocket.

To complement these single molecule biophysical studies, we also have carried out ensemble activity studies of the membrane-bound PDK1:AKT1 and PDK1:PKCα heterodimers to investigate the mechanism of phosphoactivation and the regulatory effects of heterodimer formation on kinase activity. These kinase activity measurements closely mimicked the conditions of the single molecule measurements to facilitate direct comparisons. To quantify the protein kinase activities of both PDK1 and AKT1 on the target membrane, we modified a previously described coupled PDK1-AKT1 kinase assay (33). The findings provide strong evidence that PDK1 phosphoactivates AKT1 by a *cis* activation mechanism. As predicted by the *cis* model, competitive displacement of AKT1 from the PDK1:AKT1 heterodimer with saturating PKCα or PS48 yields near complete inhibition of PDK1:AKT1 phosphoactivation reaction. Overall, the findings reveal a novel, bimodal regulatory system in which (i) moderate levels of a Ca^2+^-activated, conventional PKC can stimulate PI3K/PIP_3_ signaling in cells by phosphorylating MARCKS and releasing sequestered PIP_2_, while (ii) high levels of PKC are proposed to act as a brake on downstream PIP_3_ signaling by competitively blocking PDK1:AKT1 heterodimer formation and *cis* PDK1 phosphoactivation of AKT1. Notably, the latter inhibition of PDK1 by high levels of PKC provides a simple molecular explanation for the recent discovery that PKC acts as a tumor suppressor in cells (41–43).

## MATERIALS AND METHODS

### Reagents

Synthetic DOPIP_3_ (1,2-dioleoyl-sn-glycero-3-phospho-1’-myo-inositol-3’,4’,5’-trisphosphate), DOPC (1,2-dioleoyl-sn-glycero-3-phosphocholine), DOPS (1,2-dioleoyl-sn-glycero-3-phospho-L-serine), and LRBPE (1,2-dioleoyl-sn-glycero-3-phosphoethanolamine-N-(lissamine rhodamine B sulfonyl) were from Avanti Polar Lipids (Alabaster, AL). PMA (phorbol-12-myristate-13-acetate) was from MilliporeSigma. Alexa Fluor™ 555 C2 Maleimide and CoverWell 9 mm perfusion chambers were from Invitrogen (Carlsbad, CA). Glutathione Sepharose 4B was from GE Healthcare Bio-Sciences (Piscataway, NJ). Talon metal affinity resin was from Takara Bio (Mountain View, CA). The PKCα, PDK1 and AKT1 constructs employed were expressed in eukaryotic tissue culture cells and purified as follows. PKCα was expressed in HEK 293 cells, purified via an HA affinity tag and, where indicated, labeled on a ybbr-labeling tag by the enzymatic Sfp labeling reaction as previously described (24, 44). Alternatively, dark, unlabeled PKCα was obtained from Abcam (Cambridge UK) and exhibited behavior in single molecule studies indistinguishable from ybbr-PKCα. PDK1 and AKT1 were expressed in Sf9 cells and purified using immobilized metal affinity chromatography as previously described (45–47). Subsequently, where indicated, PDK1 and AKT1 were labeled on a ybbr-labeling tag by Sfp as previously described (23, 24, 48, 49). Figure 2C shows the locations of labeling and affinity tags in domain structures of each construct, as well as relevant phosphorylation sites. Each kinase is phosphorylated at one or more of these sites during eukaryotic cell expression as follows. PKCα isolated from tissue culture cells is known to to be phosphorylated at T497 and S657 (50, 51). PDK1 autophosphorylates its S241 and T513 sites during cell expression, and after purification when ATP and PIP_3_-containing membranes are present (32, 33). AKT1 isolated is phosphorylated at S473 during expression, but not at T308 (33, 45, 52, 53), which makes it a suitable substrate for PDK1 phosphoactivation *in vitro* (see Figs. 8,9). 5-FAM fluor-labeled Crosstide was from AnaSpec (Fremont, CA). C1 peptide was from Promega (Madison, WI). Lyophilized ATP was from Expedeon (Cambridge, MA). All other salts were from MilliporeSigma.

### Preparation of supported lipid bilayers for Single-molecule TIRFM

Previously described protocols (23–25, 48, 54, 55) were modified to generate a homogenous supported lipid bilayer on ultra-clean glass. Briefly, lipids were dissolved in chloroform (DOPS,DOPC), 20:9:1 chloroform: methanol: water (PIP_3_), or DMSO (PMA) and mixed to yield the desired composition. Lipids were dried with a roto-vap to remove solvent and rehydrated in buffer containing 25 mM HEPES, 100 mM KCl, 15 mM NaCl, 0.5 mM MgCl_2_, 20 μM EGTA, 26 μM Ca^2+^, 5 mM reduced glutathione, pH adjusted to 7.4 with 3 M KOH. The lipid mixture was vortexed for 1 minute and subjected to 3 freeze-thaw cycles in liquid N_2_ with 1-minute vortex after each thaw. After 3 freeze-thaw cycles the lipid mixtures were aliquoted into 2 mL amber screw-top tubes and snap frozen in liquid N_2_ and stored at −80 °C until use. On the morning of an experiment a lipid aliquot was thawed on ice and sonicated with a Misonix XL2020 probe sonicator with a micro tip for a total of four minutes with 0.5 second pulses followed by 0.5 second pauses. The SUVs were mixed with 1 M NaCl and deposited for 30 minutes onto an ultra-clean glass surface in the imaging chamber. After depositing the lipids were rinsed with deionized H2O and then with kinase assay buffer supplemented with 0.1 mg/mL BSA.

### Single-molecule TIRFM measurements

TIRFM experiments were carried out at 22.0 ± 0.5 °C on an objective-based TIRFM instrument, as described previously (23, 56). The instrument utilized a Nikon (Melville, NY) TE2000U inverted TIRF microscope; a Nikon Apochromat 60×, NA 1.49 TIRF oil immersion objective; and a 532 nm, diode-pumped solidstate laser (CNI-Laser model MGLIII-532-300 mW). The laser power was reduced with two N.D. filters totaling 0.8 absorbance. Sample fluorescence emerging from the 600 nm long-pass emission filter was captured by an Andor iXon Life electron-multiplying charge-coupled device camera (Oxford Instruments, Abingdon, UK).

### Single-molecule diffusion tracking and analysis

The analysis of the single molecule diffusion movies was carried out as previously described (44, 54, 55, 57). The trajectories of fluor-labeled proteins or lipids were tracked using the ImageJ plugin Particle Tracker (58). The software extracts positions and brightness for each trajectory in every frame it is present. These data are then imported into Mathematica for analysis. Particles were filtered on the basis of brightness to remove dim contaminants and excessively bright aggregates/contaminants. Additional filtering was done on the basis of displacement to remove immobile and rapidly dissociating particles. All filtering has been previously described and validated (44, 54, 55, 57). Distributions of single track diffusion constants (48), and population-averaged single molecule diffusion constants were determined as previously described (54).

### Membrane-bound densities of fluor-labeled proteins

smTIRFM protein density measurements were carried out by effectively counting the number of single molecule tracks (as determined above, after filtering) in each frame of a 10 sec movie and then averaging over the total number of frames. In practice, for convenience the frame lengths of all the filtered trajectories were summed and divided by the number of frames in the movie yielding the binding density per observation area.

### Buffer and lipid compositions

The standard assay buffer for smTIRFM and bulk activity studies was 25 mM HEPES, 100 mM KCl, 15 mM NaCl, 0.5 mM MgCl_2_, 20 μM EGTA, 26 μM Ca^2+^, 5 mM reduced glutathione, 1 mM ATP, pH 7.4 except where indicated otherwise. The standard lipid composition for these studies was 72/24/2/2 mol% DOPS/DOPC/PIP_3_/PMA except where indicated otherwise.

### Coupled PDK1-AKT1 Protein Kinase Assay

A previously described coupled PDK1:AKT1 protein kinase assay was adapted (33) for use with a fluorescent AKT1 peptide substrate. PDK1 (0.25 nM) and AKT1 (60 nM) enzymes were combined in assay buffer supplemented with 6 μM sonicated lipids (standard lipids, or 73.5/24.5/2 DOPC/DOPS/DOPIP_3_, or 75/25 DOPC/DOPS) and 100 μg/mL BSA. The reaction mixture was then mixed and incubated at 22 °C for 10 minutes. The reaction was started upon addition of ATP to a concentration of 1 mM. After 7.5 minutes of incubation at 22 °C this reaction mixture was diluted 1:4 into assay buffer supplemented with 1 mM ATP and 0.3 mg/mL 5-FAM Crosstide. This reaction was then incubated for 7.5 minutes at 22 °C and quenched by heating at 95 °C for 10 minutes. The quenched reaction mixture was cooled on ice and combined with 3 uL 80% glycerol. The product was separated from the substrate on a 0.8% agarose gel run at 200V for 25 minutes. The gels were imaged on a LAS-4000 gel imager and the band intensities calculated using ImageJ.

### PKCα Protein Kinase Assay

Enzymes were combined in assay buffer supplemented with 6 μM sonicated standard lipids, 100 μg/mL BSA, and 0.04 mg/mL C1 peptide. The reaction was initiated upon addition of ATP to 1 mM, incubated at 22 C for 20 minutes and was quenched by heating the reaction mixture to 95 C for 10 minutes. The quenched reaction mixture was cooled on ice, combined with 3 uL 80% glycerol. The product was separated from the substrate on a 0.8% agarose gel run at 200V for 25 minutes. The gels were imaged on a LAS-4000 gel imager and the band intensities calculated using imageJ.

### Statistics

Single molecule TIRFM and bulk activity experiments were carried out for at least three independently prepared trials on two or more days, where each trial was at least a triplicate. Reported n values are the number of independently prepared trials, with uncertainty reported as standard error of the mean (SEM). smTIRFM experiments were carried out with at least three movies recorded for each condition. Statistical significance was determined using a one- or two-tailed t-test as appropriate.

## RESULTS

### Preparation of full-length master kinases for single molecule studies

The present studies employ three full length master kinases purified by affinity chromatography from eukaryotic tissue culture cells: PKCα, PDK1 and AKT1. We have previously reported extensive single molecule studies of full length PKCα on supported bilayers (11, 24, 25). Here we report the first single molecule studies, to our knowledge, of fluorescently labeled, full-length PDK1 and AKT1. In experiments where a fluorescent label is required, each protein is enzymatically labeled with an Alexa Fluor probe on an N- or C-terminal ybbr labeling tag as previously described for PKCα (11, 24, 25). Figure 2B presents the domain structures and relevant phosphorylation sites of protein constructs employed.

### Measuring membrane binding, 2-dimensional diffusion, and heterodimer formation of fluor-labeled proteins on supported lipid bilayers

Single-molecule total internal reflection fluorescence microscopy (smTIRFM) enables direct measurements of the membrane docking and 2-dimensional diffusion of peripheral membrane proteins on supported lipid bilayers. Only proteins stably bound to the membrane are visualized by the method since the diffusivity of proteins in the aqueous phase is too high for imaging, while membrane-bound proteins are easily imaged because the high viscosity of the lipid bilayer reduces their diffusivities 2-3 orders of magnitude (49, 57).

Heterodimer formation between membrane-bound proteins on supported lipid bilayers can be detected by the lower diffusivity of the dimer compared to that of the individual monomers (49, 54, 57). The 2-dimensional diffusion constant D of each membrane-bound protein or complex is inversely proportional to its frictional drag, and systematic studies have shown that the total frictional drag due to multiple membrane-bound domains or oligomer subunits is additive on supported lipid bilayers (49, 54, 57). Thus, when two proteins form a heterodimer, the total frictional drag of the complex is the sum of the frictional drags of the two individual monomers, often yielding a detectable reduction of D and diffusivity. In addition, the measured monomer diffusion constants and frictional drags can be used to calculate the predicted diffusion constant of each heterodimeric complex formed between them, enabling direct comparisons between calculated and experimental heterodimer diffusion constants.

Fluor-labeled, full length PDK1, AKT1 and PKCα each recognize and bind to specific lipids on a target membrane surface. PDK1 and AKT1 possess a PIP_3_-specific PH domain and exhibit stable binding to a supported lipid bilayer, but only when the bilayer contains target lipid PIP_3_, as demonstrated in **Figure 3**. PKCα possesses multiple membrane targeting domains, including the C2 domain that recognizes PS, which is required for effective membrane docking as shown in Figure 3, as well as C1A and C1B domains that recognize PMA. To confirm the target lipid specificity of each protein, PDK1 (20 pM) or AKT1 (50 pM) or PKCα (50 pM) labeled with Alexa-Fluor 555 was imaged on bilayers containing or lacking its key target lipid (Figure 3). The density of fluor-labeled kinase molecules per field was determined by averaging the number of freely-diffusing fluor-kinase molecules detected in each video frame over all frames of the movie (algorithm in Methods). Figure 3 shows that the average density of each labeled protein decreased 4-fold to 100-fold when its target lipid was omitted from the supported lipid bilayer.

**Figure 3.**
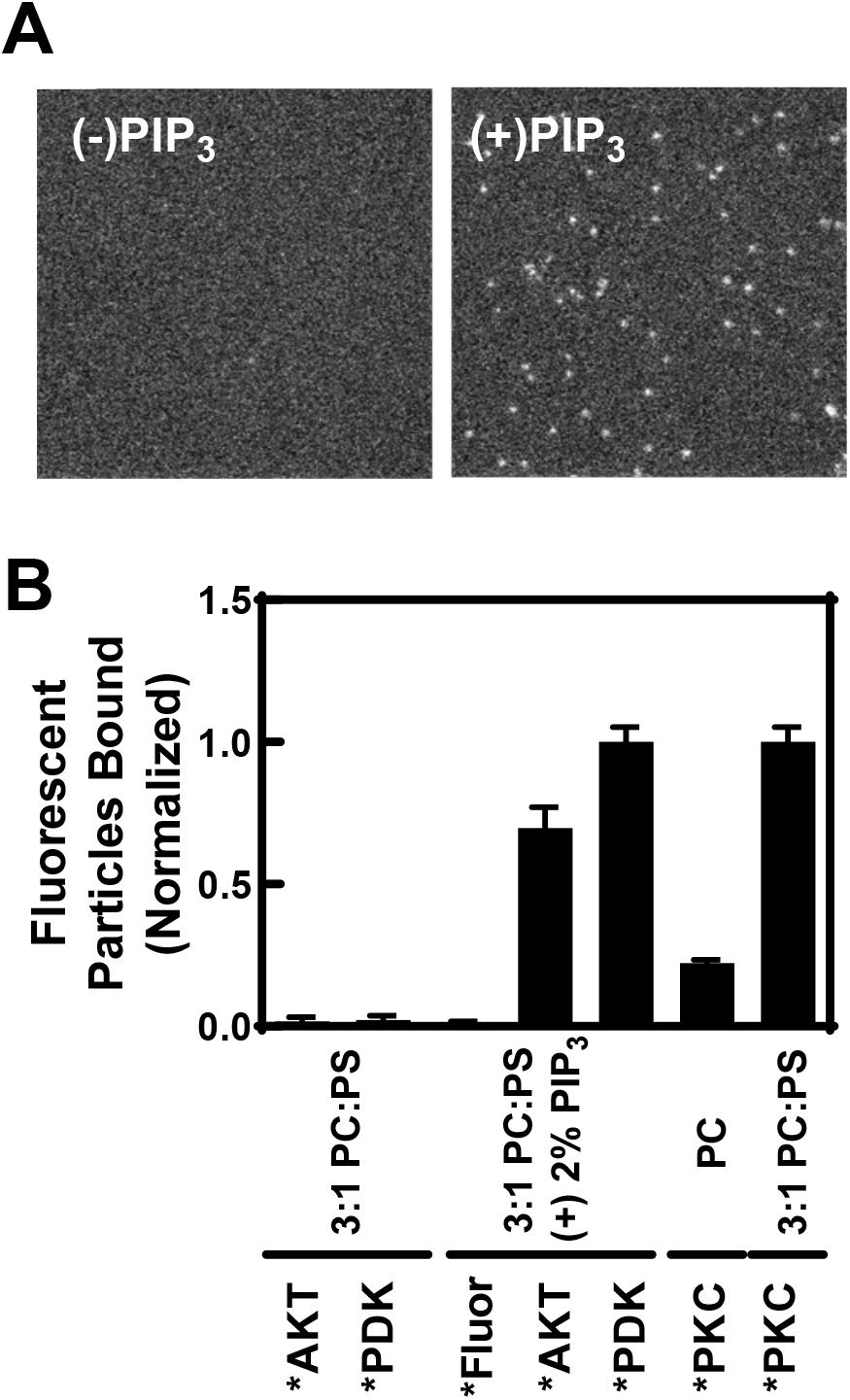
Single molecule TIRFM quantitation of protein kinase binding to supported lipid bilayers lacking and containing target lipid. **(A)** Fluor-labeled PDK1 was imaged on bilayers composed of 75/25 mol% DOPC/DOPS (left) and 73.5/24.5/2 mol% DOPC/DOPS/DOPIP_3_ (right). **(B)** Membrane-bound density of fluor-labeled PDK1 and AKT1 measured on bilayers containing DOPC/DOPS 75/25 mol% and 73.5/24.5/2 mol% DOPE/DOPS/DOPIP_3_, showing that density increases ~100 fold (p < 0.001) when PIP_3_ is present. Membrane-bound density of fluor-labeled PKCα was measured on bilayers containing pure DOPC 100 mol% and DOPC/DOPS 75/25 mol%, showing that density increases ~4-fold (p < 0.001) in the presence of PS. (PKCα data obtained as previously described (44)). Asterisk indicates AF555 fluor-labeled protein. The control *Fluor is an equivalent concentration of free AF555-glutathione, testing whether free fluor contributes to signal. Each average density was obtained from at least four independently prepared samples (n≥4) measured on at least two different days, where each sample was imaged in at least three different movies.

**Figure 4** illustrates the effect of heterodimer formation on 2-D diffusion. Representative single-molecule position trajectories are plotted for PDK1 monomers and PDK1 heterodimers with AKT1 and PKCα. Both the step lengths (the distance traveled between 20 msec frames) and the total displacements observed for PDK1 monomer trajectories (Fig. 4A) decrease in the presence of saturating AKT1 (Fig. 4A) or PKCα (Fig. 4A). Thus, PDK1 forms heterodimers with both AGC kinases. The average diffusion coefficients of the species in Figure 4A are compared in Figure 4B, confirming that the diffusion coefficient of fluor-PDK1 is significantly reduced by the addition of saturating dark AKT1 (p < 0.001) or PKCα (p < 0.001) due to formation of PDK1:AKT1 or PDK1:PKC heterodimers, respectively, with the concomitant increase in frictional drag against the bilayer.

**Figure 4.**
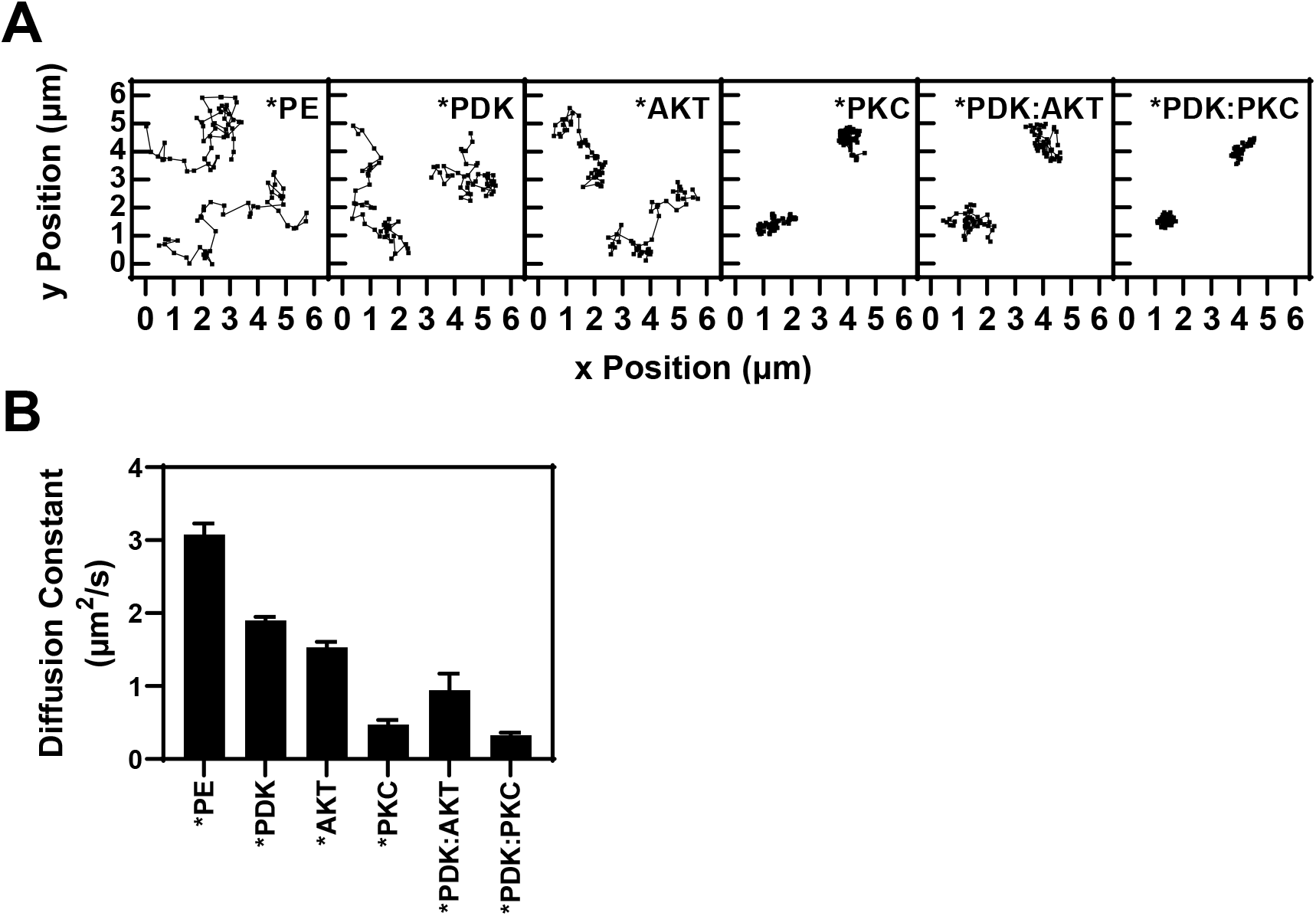
Single molecule TIRFM analysis comparing the 2-D diffusion of membranebound protein kinases and their heterodimers. **(A)** Plots of representative single molecule 2-D diffusion trajectories on supported lipid bilayers, with individual diffusion steps defined by the 20 ms video frame exposure time. From left to right: fluor-labeled lipid LRB-PE, fluor-labeled PDK1, fluor-labeled AKT1, fluor-labeled PKCα (generated as described in (44)), fluor-labeled PDK1 with saturating dark AKT1, fluor-labeled PDK1 with saturating dark PKCα. **(B)** The average diffusion coefficient of each species. The D of PKCα is significantly smaller than those of PDK1 (p < 0.001) and AKT1 (p < 0.001) due to its greater, multi-domain frictional drag against the bilayer (Figure 2 and (11)). The D of PDK1 is significantly decreased by the binding of AKT1 (p < 0.001) or of PKCα (p < 0.001) to form a heterodimer. Each average diffusion constant was obtained from at least six independently prepared samples (n≥6) measured on at least three different days, where each sample was imaged in at least three movies.

### Equilibrium affinity and PIF interaction of the membrane-bound PDK1:AKT1 heterodimer

To measure the affinities of membrane-bound heterodimers, fluor-labeled, membrane-bound PDK1 was titrated with its dark AGC kinase binding partner, while measuring the average diffusion coefficient of the fluor-PDK1 population by smTIRFM analysis. Previous studies have shown that in such a titration, the average 2-D diffusion constant of the fluorescent protein population is the average of the monomer and heterodimer diffusion constants, weighted by their relative proportions (49). The titration begins with the D value of the fluorescent monomer and asymptotically approaches the D value of the heterodimer, which is smaller due to the additive frictional drags of the two binding partners against the bilayer. Plotting the average diffusion constant versus the bulk concentration of dark AGC kinase thus yields a simple binding curve with the K_1/2_ revealing the bulk AGC kinase concentration needed to convert half of the membrane bound PDK1 population to heterodimers. The resulting K_1/2_ can be compared with the bulk AGC kinase concentration in cells to help ascertain the physiological relevance of the observed heterodimer.

**Figure 5** begins by showing distributions of single track diffusion constants for fluor-PDK1 as dark AKT1 is added (Figure 5A). As expected, the diffusion constant distribution observed for individual fluor-PDK1 molecules shifts to lower D as [AKT1] increases and each fluor-PDK1 monomer spends a greater fraction of its time bound to AKT1. Figure 5B plots the mean fluor-PDK1 diffusion constant, averaged over the entire fluor-PDK1 population, versus the bulk dark AKT1 concentration, yielding a simple binding curve with a K_1/2_ of 30 ± 8 nM (Figure 5B). The asymptote at high AKT1 concentration yields the diffusion coefficient of the PDK1:AKT1 heterodimer, D = 0.94 ± 0.06 μm^2^/sec. Figure 5C shows this experimental heterodimer diffusion constant is significantly smaller (p < 0.001) than the diffusion constants of the PDK1 monomer (D = 1.99 ± 0.12 μm^2^/sec) and of the AKT1 monomer (D = 1.53 ± 0.07 μm^2^/sec). Further, the measured heterodimer diffusion constant is the same (Figure 5C), within error, as the predicted value (D = 0.86 ± 0.07 μm^2^/sec) calculated by summing the frictional drags of the PDK1 and AKT1 monomers owing to the additivity of frictional drags on supported lipid bilayers (57).

**Figure 5.**
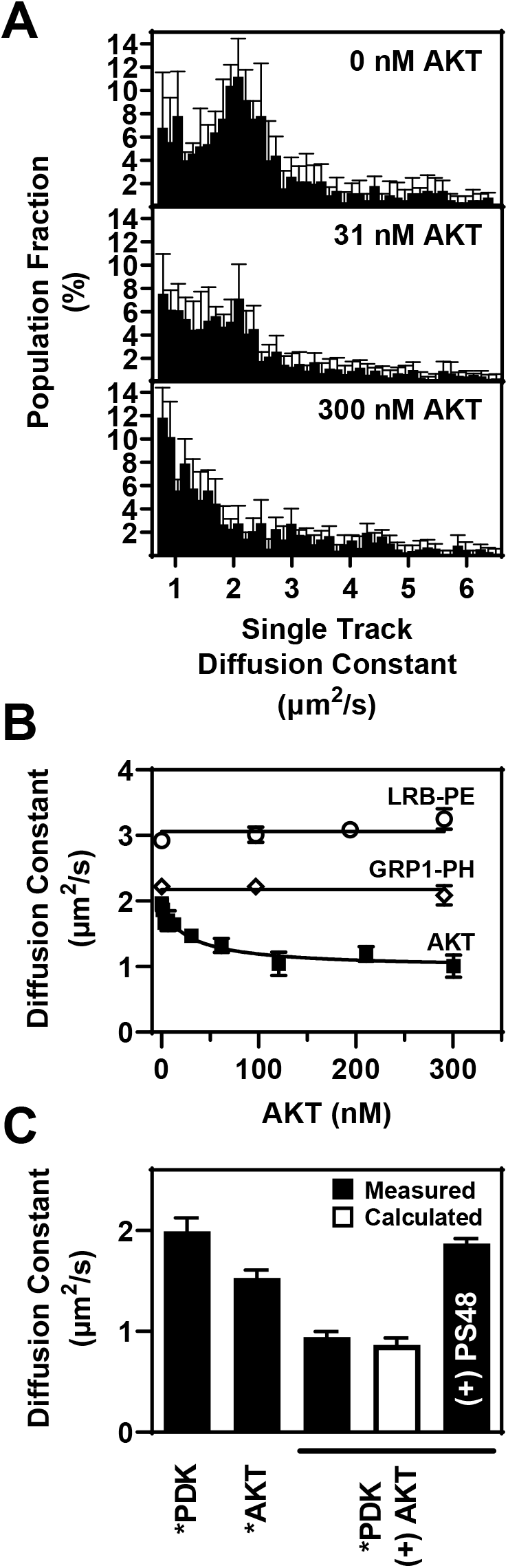
Single molecule TIRFM quantitation of AKT1 binding to PDK1. **(A)** The distribution of fluor-PDK1 single molecule diffusion coefficients is plotted without AKT1 (top panel), and at two dark AKT1 concentrations (lower panels) that shift the distribution to lower diffusivity. Population fractions are from four independent samples, each imaged three times (n=4). The low diffusivity subpopulation in the absence of AKT may represent PDK1 homodimers. **(B)** Titration monitoring the average diffusion constant of fluor PDK1 as dark AKT1 is added in increasing concentration. The resulting best-fit K_1/2_ for AKT1 binding to PDK1 is 30 ± 9 nM. The diffusion constants o f negative controls fluorlipid LRB-PE and fluor-protein GRP1PH are relatively independent of AKT1 addition, indicating AKT1 does not alter general bilayer diffusion. Average diffusion constants are from four independent samples, each imaged three times (n=4). **(C)** Comparison of the indicated average diffusion constants, indicating that addition of saturating (300 nM) dark AKT1 converts the PDK1 population to heterodimers, yielding an average diffusion constant significantly lower (p < 0.001) than that of PDK1 monomer alone, but within error of that predicted for the heterodimer based on the additive frictional drags of the monomeric kinases. The small molecule PS48 was found to disrupt the PDK1:AKT1 heterodimer and increase (p < 0.001) the average fluor-PDK1 diffusion constant nearly to its monomer value, indicating that PS48 competitively displaces AKT1 from the heterodimer. Each average diffusion constant was obtained from four to six independently prepared samples (n≥4) measured on at least two different days, where each sample was imaged in at least three movies.

Controls were carried out to ascertain whether titration of the membrane system with dark AKT1 altered the diffusion of lipids or non-interacting proteins on the membrane surface. The diffusion of a fluorescent lipid (LRB-PE) in the supported bilayer, or fluor-labeled GRP1-PH domain bound to PIP_3_ in the bilayer, was monitored while titrating in dark AKT1 (Figure 5B). The increasing AKT1 concentration had no detectable effect on the diffusion coefficient of LRB-PE or GRP1-PH, indicating that the effect of AKT1 on the PDK1 diffusion coefficient is due to a specific PDK1-AKT1 binding interaction, rather than a non-specific decrease in fluidity of the supported lipid bilayer.

To ascertain whether the PDK1:AKT1 heterodimer is stabilized by a PIF interaction between the AKT1 PIF motif and the PDK1 PIF pocket, the effect of the PIF pocket-binding drug PS48 on the heterodimer was examined. Figure 5C shows that addition of saturating PS48 does disrupt the PDK1:AKT1 heterodimer and significantly increases (p < 0.001) the average diffusion constant to nearly the same level observed for the PDK1 monomer. These findings provide direct evidence that the PDK1:AKT1 heterodimer is stabilized by a PIF interaction.

### Equilibrium affinity and PIF interaction of the membrane-bound PDK1:PKCα heterodimer

The same approach was used to determine the equilibrium affinity of the PDK1:PKCα heterodimer. **Figure 6A** shows that the diffusion constant distribution observed for individual fluor-PDK1 proteins shifts to lower D as dark PKCα is added and causes a given fluor-PDK1 monomer to spend a greater fraction of its time bound to PKCα. Figure 6B displays the titration of the fluor-PDK1 diffusion constant, averaged over the imaged population, with dark PKCα. Interestingly, the average PDK1 diffusion constant was unaltered until the dark PKCα concentration exceeded 2.8 nM, possibly due to PKCα sticking to surfaces of the imaging chamber until those surfaces are blocked. Fitting the rest of the titration to a single-site binding equation yields K_1/2_ = 3.8 ± 0.4 nM as shown in Figure 6B. Figure 6C indicates that the asymptotic PDK1:PKCα heterodimer diffusion constant at saturating PKCα (D = 0.33 ± 0.03 μm^2^/sec) is significantly smaller (p < 0.001) than the diffusion constant of the PDK1 monomer (D = 1.90 ± 0.04 μm^2^/sec), and is indistinguishable from the predicted heterodimer diffusion constant (D = 0.38 ± 0.14 μm^2^/sec). The large frictional drag of PKCα arises from its multiple, bilayer-embedded lipid binding domains (Figure 2) and accounts for the low D of both the PKCα monomer and the PDK1:PKCα heterodimer. The ability of PKCα to decrease PDK1 diffusivity is due to a specific PDK1:PKCα interaction, and not due to a non-specific decrease in fluidity of the supported lipid bilayer, since titration with dark PKCα has no effect on the diffusion constants of fluor-labeled lipid or non-interacting GRP1-PH domain bound to PIP_3_ (Figure 6B)

**Figure 6.**
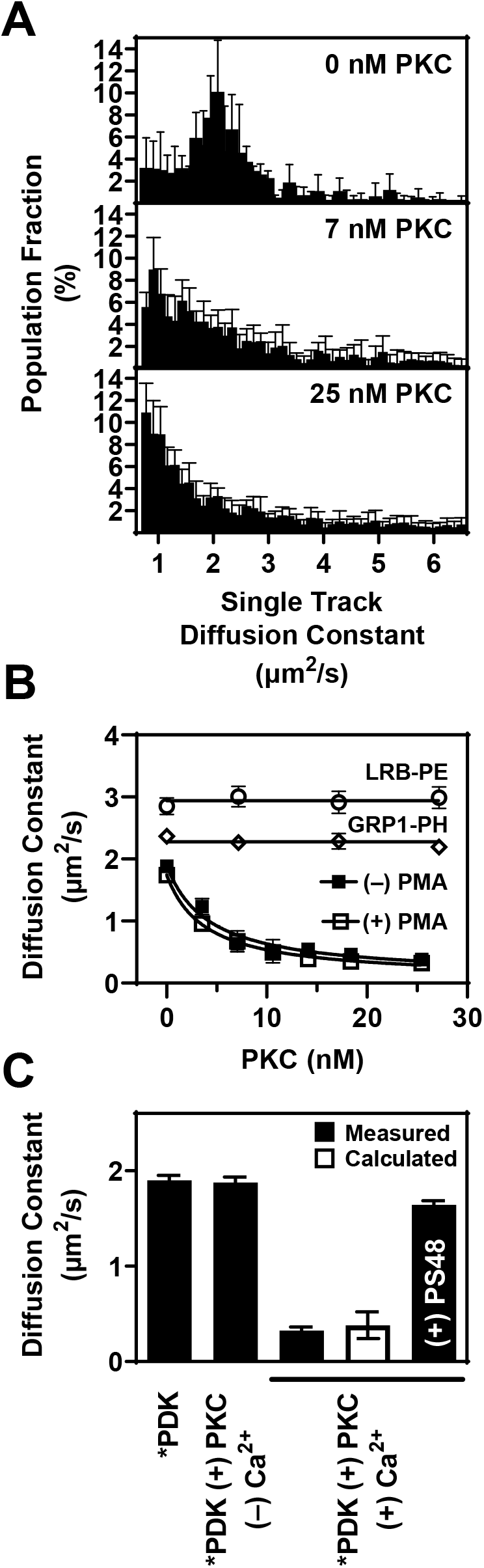
Single molecule TIRFM quantitation of PKCα binding to PDK1. **(A)** The distribution of fluor-PDK1 single molecule diffusion coefficients is plotted without PKCα (top panel), and at two dark PKCα concentrations (lower panels). Population fractions are from four independent samples, each imaged three times (n=4). **(B)** Titration monitoring the average diffusion constant of fluor-PDK1 as dark PKCα is added in increasing concentration. The resulting best-fit K_1/2_ for PKCα binding to PDK1 is 3.8 ± 0.4 nM with PMA, and 4.2 ± 0.7 nM without PMA. The diffusion constants of two negative controls, fluor-lipid LRB-PE and fluor-protein GRP1PH, are relatively independent of PKCα addition, indicating PKCα does not alter general bilayer diffusion. Diffusion constants are from four independent samples, each imaged three times (n=4). (At the lowest concentrations PKCα is recruited to surfaces and no fluor-PDK1 diffusion change is observed until PKCα reaches 2.8 nM; thus, when fitting the data, the lowest concentration points were excluded and the y-axis was shifted to transform 2.8 nM PKCα to 0 nM). **(C)** Comparison of the indicated average diffusion constants, indicating that when Ca^2+^ is present, addition of saturating (25 nM) dark PKC α converts the PDK1 population to heterodimers, yielding a significantly decreased (p < 0.001) average heterodimer diffusion constant indistinguishable from that predicted for the heterodimer based on the additive frictional drags of the monomeric kinases. The small molecule PS48 was found to disrupt the PDK1:PKCα heterodimer and increase (p < 0.001) the average fluor-PDK1 diffusion constant nearly back to its monomer value, indicating that PS48 competitively displaces PKCα from the heterodimer. Diffusion constants are from four or five independent samples (n=4-5), each imaged three times on at least two different days.

Additional experiments investigated the effects of PKCα activators on PDK1:PKCα heterodimer formation. The same K_1/2_ for PKCα binding to PDK1 was observed, within error, when the lipid bilayer contained or lacked the PKCα activating lipid, phorbol 12-myristate 13-acetate (PMA). It follows that the affinity of the PDK1:PKCα heterodimer is independent of the PKCα activation state (Figure 6A). Further, the PKCα-triggered loss of PDK1 diffusivity was no longer observed when Ca^2+^ is removed (Figure 6B), as expected since PKCα requires Ca^2+^ for membrane binding. This finding is consistent with a loss of PDK1:PKCα heterodimers in the absence of Ca^2+^.

To ascertain whether the PDK1:PKCα heterodimer is stabilized by a PIF interaction between the PKCα PIF motif and the PDK1 PIF pocket, the effect of the PIF pocket-binding drug PS48 on the heterodimer was examined. Figure 6C shows that addition of saturating PS48 does disrupt the PDK1: PKCα heterodimer and significantly increases (p < 0.001) the average diffusion constant towards the original level observed for the PDK1 monomer. These findings directly show that the PDK1:PKCα heterodimer is stabilized by a PIF interaction and that the heterodimer is dissociated by a competitive inhibitor that targets the PIF pocket.

### PKCα competitively displaces AKT1 from the PDK1:AKT1 heterodimer

Given the significantly higher affinity of the PDK1:PKCα heterodimer relative to the PDK1:AKT1 heterodimer, as well as the stabilization of both heterodimers by a PIF interaction with the PDK1 PIF pocket, we asked whether PKCα can competitively displace AKT1 from the PDK1:AKT1 heterodimer. **Figure 7A** shows that addition of saturating PKCα to the fluor-PDK1:AKT1 heterodimer causes the average fluor-PDK1 diffusion constant to drop to the same value, within error, as observed for the fluor-PDK1:PKCα heterodimer. Moreover, Figure 7B shows addition of saturating PKCα to the PDK1:fluor-AKT1 heterodimer causes the average diffusion constant of fluor-AKT1 to increase to nearly the same level as the AKT1 monomer. These findings indicate that PKCα and AKT1 bind competitively to the PIF pocket on PDK1, and demonstrate that the higher affinity PKCα can effectively displace AKT1 from the PDK1:AKT1 heterodimer to yield the more stable PDK1:PKCα heterodimer and free AKT1.

**Figure 7.**
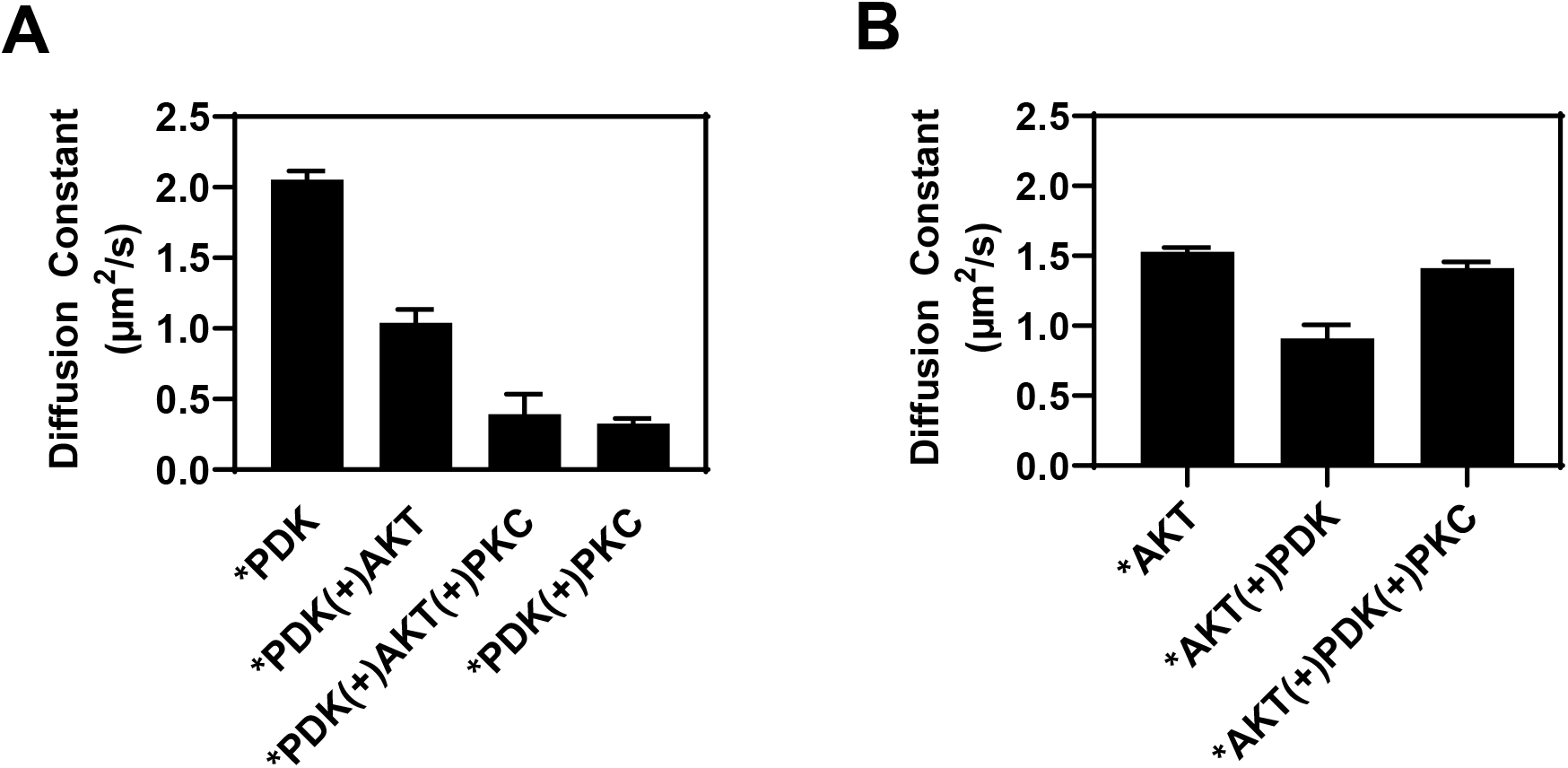
AKT1 and PKCα compete for binding to PDK1. **(A)** Fluor-PDK1 diffusion coefficient is monitored with additions of dark AKT1 (60 nM) and a subsequent addition of dark PKCα (31 nM). Labeled PDK1 with dark PKC is shown as a reference. Addition of dark AKT1 significantly reduces the average fluor-PDK1 diffusion constant (p < 0.001) upon formation of the PDK1:AKT1 heterodimer, while subsequent addition of dark PKCα displaces AKT1 and further decreases D (p < 0.001) to nearly the same level observed for the PDK1:PKCα heterodimer. Diffusion constants are from three independent samples, each imaged three times on two different days (n=3). **(B)** Fluor-AKT1 diffusion coefficient is monitored with addition of dark PDK1 (80 nM) and then subsequent addition of dark PKCα (31 nM). The fluor-AKT1 D decreases upon addition of dark PDK1 (p < 0.001), then further addition of dark PKCα increases D (p = 0.004) nearly back to its fluor-AKT1 monomer level as the PKCα displaces fluor-AKT1 and forms PDK1:PKCα heterodimers. Diffusion constants are from six independent samples (n=6), each imaged three times on two different days.

### Regulatory effects of PDK1:AKT1 or PDK1:PKCα heterodimer formation on the individual kinase activities of PDK1, AKT1 and PKCα

We next established bulk activity measurements utilizing essentially the same conditions (buffer, ionic composition, membrane lipid composition and net accessible lipid, and protein concentrations) as employed in the single molecule studies to facilitate direct comparisons. As previously noted by others (33), our AKT1 enzyme was not yet phosphoactivated and exhibited negligible kinase activity prior to PDK1 phosphoactivation. To analyze the protein kinase activities of both PDK1 and AKT1 on their target membrane, we modified a previously described coupled PDK1-AKT1 kinase assay (33) as illustrated in **Figure 8**.

**Figure 8.**
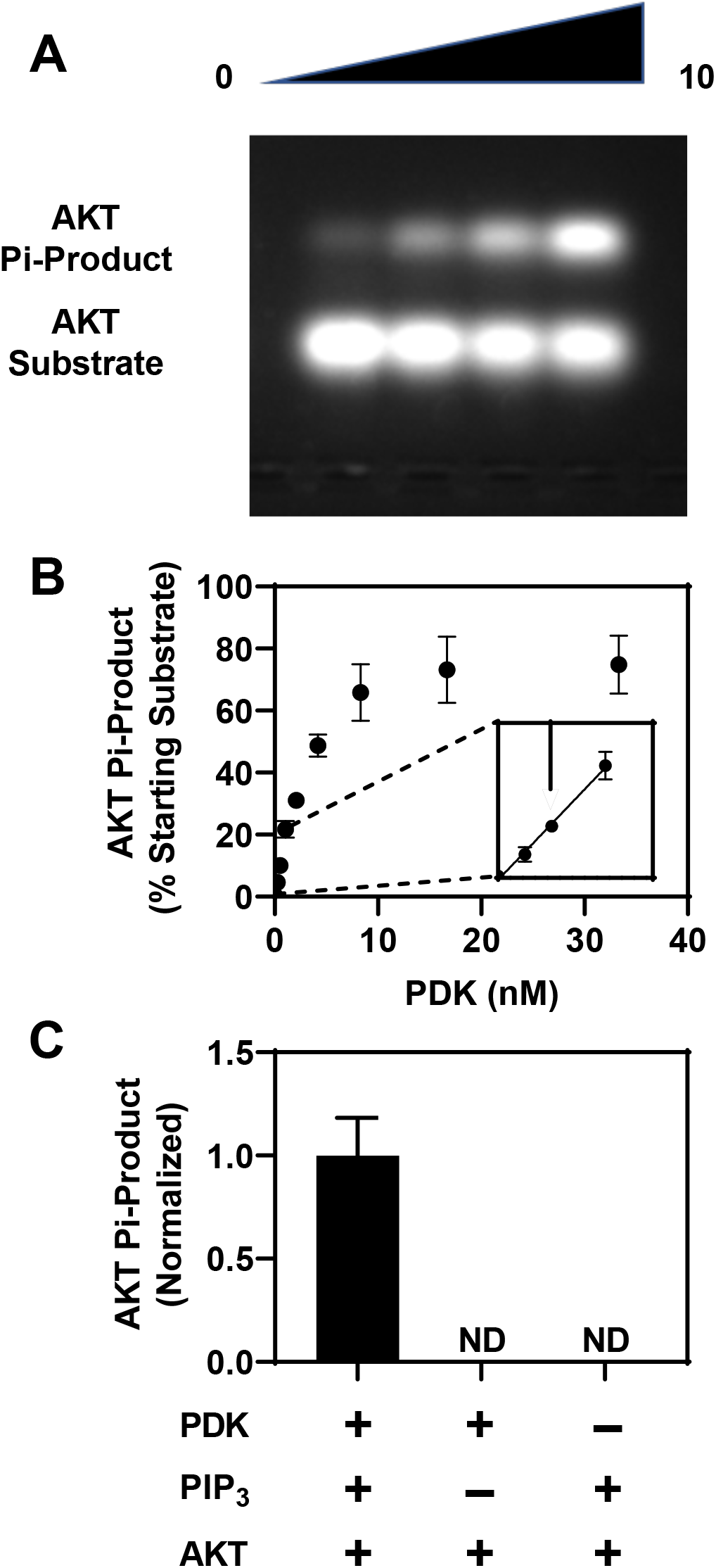
Coupled PDK1/AKT1 protein kinase assay, modified from a previous coupled assay (33). In this two-phase assay, PDK1 and AKT1 are first incubated with target PIP_3_ membranes and ATP to drive PDK1 phosphoactivation of AKT1. Then AKT1 fluorescent substrate is added and a second incubation is carried out to measure AKT1 protein kinase activity. **(A)** AKT1 substrate peptide is separated from phosphorylated product peptide (Pi-product) on an agarose gel and both bands are integrated to quantify the fraction of substrate converted to product. **(B)** Titration of PDK1 in the coupled assay to find the range in which AKT1 activity increases linearly with PDK1 concentration. The arrow in the expanded inset shows the PDK1 concentration used in further experiments. Activity data are from four independent reactions on two different days (n=4). **(C)** Controls showing that no AKT1 activity is detected in the absence of phosphoactivation by PDK1, nor in the absence of PIP_3_. ND indicates that AKT1 Pi-product was not detected. Activity data are from three independent reactions (n=3) measured on two different days, each determined in triplicate.

Using the coupled PDK1-AKT1 kinase assay, we first asked if the PDK1 phosphoactivation of AKT1 occurs via a *cis* or *trans* reaction. In both models, the phosphoactivation reaction is catalyzed by an activated PDK1 molecule in a heterodimer with its PIF pocket occupied by the PIF motif of its AGC kinase binding partner as needed for full PDK1 activation (28, 59–61). In the *cis* model, which is more widely presented in the literature (62–64), the activated PDK1 in a PKD1:AKT1 heterodimer phosphorylates the AKT1 molecule within the same heterodimer. In the *trans* model, activated PDK1 phosphorylates an AKT1 molecule outside its own heterodimer; either a freely diffusing AKT1 monomer or an AKT1 molecule in a different heterodimer. To our knowledge, the *cis* and *trans* models have not yet been experimentally resolved yet they make simple, opposing predictions about the effect of PKCα addition on the PDK1-catalyzed phosphoactivation of AKT1 in the coupled PDK1-AKT1 kinase assay. The *cis* model predicts that of addition of saturating PKCα (or the PIF pocket binding drug PS48) will block PDK1 phosphoactivation of AKT1, because PKCα (or PS48) binding to the PDK1 PIF pocket will displace AKT1 from all PDK1:AKT1 heterodimers and prevent the proposed *cis* phosphoactivation reaction within the same heterodimer. In contrast the *trans* model predicts that addition of saturating PKCα (or PS48) to the coupled kinase reaction may have little or no effect on the PDK1 phosphoactivation of AKT1, because PDK1 will remain activated by PKCα (or PS48) occupancy of its PIF pocket and will retain the ability to phosphorylate freely diffusing AKT1 molecules via random collisions during their mutual 2D diffusion on the membrane surface.

**Figure 9** provides strong evidence supporting the *cis* model in which PDK1 phosphoactivates AKT1 bound within the same PDK1:AKT1 heterodimer. Thus, addition of excess PKCα, or the PIF pocket drug PS48, strongly inhibits the activation of AKT1 by PDK1 in the coupled PDK1-AKT1 kinase assay (Figure 9A,B). Figure 9C further quantifies the PKCα inhibition of the coupled PDK1-AKT1 kinase assay by titrating the assay with increasing PKCα concentrations on supported bilayers containing or lacking the activating lipid PMA. The resulting, best-fit apparent inhibition constants are K_i_(PKCα) = 17 ± 2 nM and 21 ± 4 nM respectively, indicating again that PMA does not detectably modulate the PDK1: PKCα interaction. Notably, assuming simple competitive binding of PKCα and AKT1 to the same PIF pocket on PDK1, the apparent K_i_ for competitive PKCα binding to the PDK1:AKT1 heterodimer can be written as K_i_(PKCα) = K_1/2_(PKCα) {1 + [AKT1]/K1/_2_(AKT1)}. The K_1/2_(PKCα) and K_1/2_(AKT1) values for PKCα and AKT1 binding to PDK1, respectively, were determined independently by smTIRFM analysis (Figures 5, 6) while the AKT1 concentration in the coupled kinase assay is known. Thus one can ask whether the K_i_(PKCα) measured by PKCα titration in the bulk coupled kinase assay (Figure 9) is consistent with the predicted K_i_(PKCα) based on the single molecule K_1/2_(PKCα) and K_1/2_(AKT1) heterodimer affinities (Figures 5, 6). Indeed, the predicted K_i_(PKCα) values (11 ± 4 nM and 13 ± 4 nM for measurements with and without PMA, respectively) are close to the corresponding K_i_(PKCα) values measured in the bulk coupled kinase assay. This excellent agreement provides strong support for the *cis* model for PDK1 phosphoactivation of AKT1, and for competitive binding of AKT1 and PKCα to PDK1. Such agreement also indicates that the single molecule and ensemble methods are analyzing the same 2-D heterodimer formation reactions on the target membrane surface, as ensured by the closely matched membrane, protein and buffer conditions of the two methods. These findings demonstrate the ability of PKCα to act as a strong inhibitor of PDK1 protein kinase activity by competitively displacing its AGC kinase substrate from the active PDK1 heterodimer.

**Figure 9.**
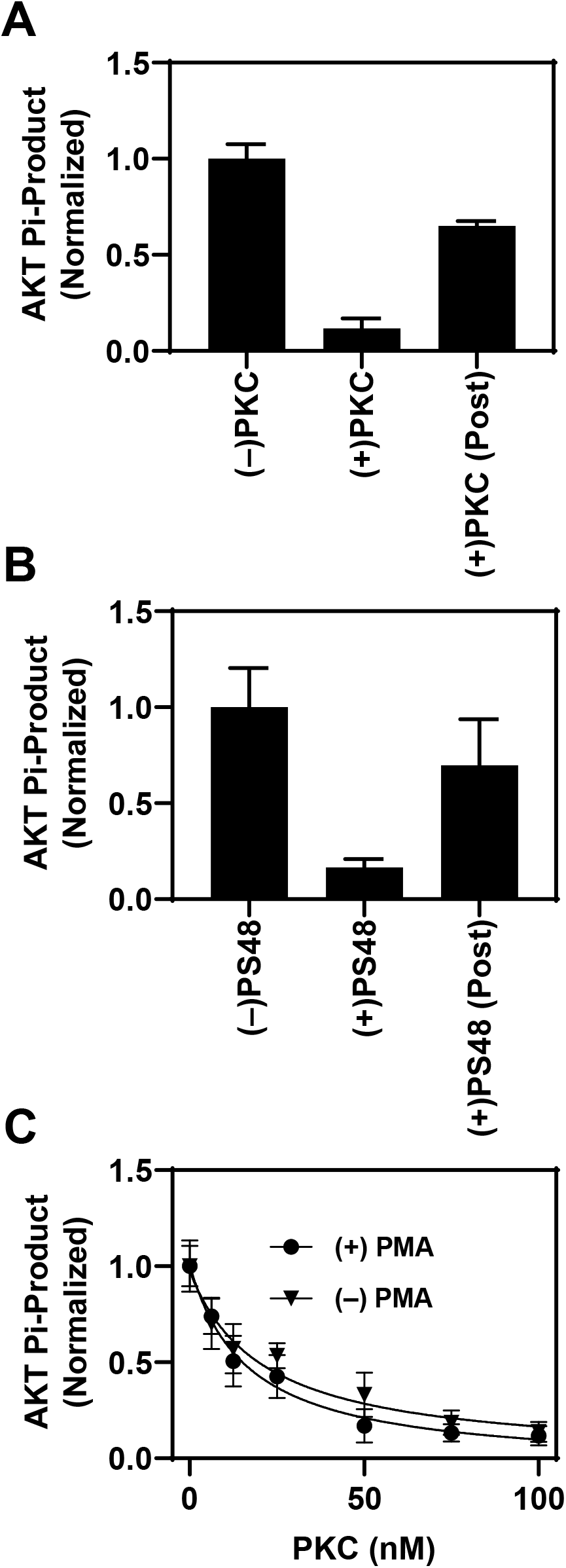
PKCα inhibits PDK1 phosphoactivation of AKT1 by competitively displacing AKT1 from PDK1 during the PDK1/AKT1 coupled kinase assay. **(A)** Addition of saturating PKCα (100 nM) to displace AKT1 from PDK1:AKT1 heterodimers during the PDK1 phosphoactivation phase (Fig. 8) prevents PDK1 phosphoactivation of AKT1, yielding significant loss (p < 0.001) of AKT1 kinase activity. In contrast, addition of the same PKCα following the PDK1 phospho-activation phase yields significantly less inhibition (p < 0.001) of AKT1 kinase activity. Data are from three independent reactions on different days, with each reaction done in triplicate (n=3). **(B)** Similar results are obtained when the PIF pocket inhibitor PS48 is added during the PDK1 phosphoactivation phase to competitively displace AKT1 from the PDK1:AKT1 heterodimer, yielding significant loss of AKT1 activity (p = 0.016). Again, significantly less inhibition is observed when PS48 is added after that phase (p = 0.047). Data are from three independent reactions with each reaction done in triplicate (n=3), on three days. **(C)** Titration of the PKCα concentration added at the beginning of the PDK1 phosphoactivation phase, yielding a K_i_ of 17 ± 2 nM. This K_i_ is quantitatively consistent competitive binding of AKT1 and PKCα to PDK1 (see text). Again, removal of PMA has no effect on K_i_ (21 ± 4 nM), providing additional evidence that the PKCα C1 domains have no measurable impact on PDK1: PKCα heterodimer stability. Data are from four (without PMA) or six (with PMA) independent reactions (n=4,6) carried out on at least four days, with each reaction done in quadruplicate.

Another relevant question is whether PDK1 detectably modulates PKCα activity in the PDK1:PKCα heterodimer. **Figure 10** shows that excess PDK1 little or no effect on the ability of PKCα to phosphorylate a soluble peptide substrate, while the PKCα inhibitor Go6976 largely blocks the same reaction. It follows that the PIF interaction within the PDK1:PKCα heterodimer has little allosteric and/or steric impact on PKCα phosphorylation of a peptide substrate, although we cannot rule out a possible larger effect on the phosphorylation of full length protein substrates where stronger steric constraints may pertain.

**Figure 10.**
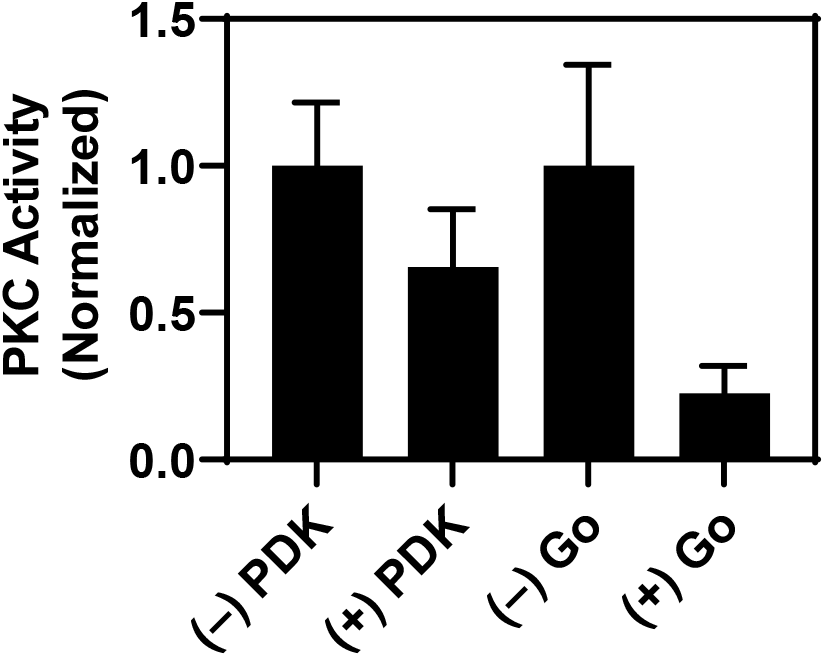
PDK1 binding to PKCα has a modest effect on PKCα protein kinase activity. Shown is a normalized protein kinase assay wherein PKCα phosphorylates a substrate p eptide. Addition of saturating (150 nM) PDK1 to drive PKCα into PDK1:PKCα heterodimers has little or no effect on the PKCα kinase activity (p > 0.35), while addition of the PKCα inhibitor Gö6976 decreases kinase activity 5-fold (p = 0.05). All activity data are from three or four independent reactions (n=3,4), each done in triplicate on two different days.

Overall, the activity studies show that when simultaneous Ca^2+^ and PIP_3_ signals recruit PKCα, PDK1 and AKT1 to the same membrane region, the high affinity PDK1:PKCα interaction may lead to competitive displacement of AKT1 from PDK1, thereby inhibiting PDK1 phosphoactivation of AKT1. Such inhibition could have important regulatory consequences for cellular signaling.

## DISCUSSION

While previous studies have implied the existence of transient PDK1:AKT1 and/or PDK1:PKCα heterodimers, the present study is the first to directly detect and analyze these heterodimers on target membrane surfaces with respect to their biophysical parameters and regulatory functions. We find that PDK1 forms stable heterodimers with both AKT1 and PKCα on target membranes possessing key plasma membrane lipids under near-physiological conditions. Single molecule detection and analysis of the membrane-bound heterodimers yields equilibrium K_1/2_ values for heterodimer formation (K_1/2_ = 30 ± 9 nM and K_1/2_ = 4.2 ± 0.7 nM for PDK1:AKT1 and PDK1:PKCα, respectively) that are near the cellular concentrations of the three kinases (for example, in HeLa cells [PDK1] ~ 42 nM, [AKT1] ~ 44 nM, and [PKCα] ~ 110 nM) (65). Thus, the observed affinities are consistent with the expected importance of these heterodimers in cell signaling pathways. The findings directly show that PDK1 forms both PDK1:AKT1 and PDK1:PKCα heterodimers through a PIF interaction between the PIF motif of the substrate kinase and the PIF pocket of PDK1. As a result, AKT1 and PKCα are observed to compete with each other for binding to the PDK1 PIF pocket, while the PIF pocket competitive inhibitor PS48 disrupts both heterodimers. Notably, the equilibrium affinity of the PDK1:PKCα heterodimer is 8-fold higher than that of the PDK1:PKCα heterodimer, ensuring that PKCα can outcompete AKT1 for PDK1 binding in many circumstances.

Ensemble kinase activity measurements were carried out under conditions closely matched to the single molecule studies to facilitate direct comparisons. One question addressed was whether PDK1 phosphoactivates AKT1 via a *cis* or *trans* mechanism. It was clear from past studies that PDK1 phosphorylates peptide substrates via a *cis* mechanism at much higher rates than it does through a *trans* mechanism, although it was unclear to us if that would remain true for full length protein substrates. We found that the *cis* mechanism pertains for full length PDK1 phosphoactivation of full length AKT1, since such phosphoactivation is lost when PDK1:AKT1 heterodimers are disrupted by competitive displacement of AKT1 from the PIF pocket by PS48 or PKCα, even though occupancy of the PIF pocket by PS48 or the PKCα PIF motif is known to activate the PDK1 kinase domain and would continue to support *trans* phosphorylation of colliding AKT1 monomers. These findings indicate that the preponderance of AKT1 substrate molecules are phosphoactivated while simultaneously bound to both the PDK1 PIF-pocket and the PDK1 active site under physiological conditions. Such simultaneous binding of the same AKT1 molecule to two clefts on opposite sides of the PDK1 kinase domain is presumably made possible by the largely unstructured AGC domain providing 60 residues and, we propose, a long, flexible tether between the AKT1 protein kinase domain and the PIF motif (Figure 2).

We also examined the ability of the PDK1, AKT1 and PKCα kinases to regulate the kinase activities of their binding partners in the PDK1:AKT1 and PDK1:PKCα heterodimers. In principle, such regulation could generate positive or negative feedback in pathways especially when simultaneous, strong Ca^2+^ and PIP_3_ signals cause high membrane densities of the three kinases and buildup of the heterodimers. Most importantly, the present findings demonstrate that PDK1:PKCα heterodimer formation can dramatically inhibit the predominantly *cis* PDK1 phosphoactivation of AKT by competitively displacing AKT1 from PDK1 under physiological conditions. The effectiveness of PKCα as a competitive inhibitor of PDK1-AKT1 phosphoactivation stems from both the 8-fold higher affinity of PKCα for PDK1, as well as the ~ 2.5-fold higher concentration of PKCα in HeLa cells (see above). In cellular settings, the novel role of PKCα as an inhibitor of PDK1-AKT1 phosphoactivation will be most important when the membrane density of Ca^2+^-activated PKCα is high, for example during a strong Ca^2+^ signal that drives global PKCα docking to plasma membrane (66, 67), or at the leading-edge membrane of polarized leukocytes where kinetically stable, local Ca^2+^ and DAG signals are known to recruit high local densities of PKCα (11, 25, 68–72). We hypothesize that under these conditions formation of the PDK1:PKCα heterodimer downregulates PDK1 phosphoactivation of AKT1 and other AGC kinases, thereby preventing runaway PIP_3_ pathway activation. In addition, while mature PKCα is already phosphorylated by PDK1 and its protein kinase activity is relatively independent of PDK1 binding (Figure 10), incorporation of mature PKCα into PDK1:PKCα heterodimers could inhibit further PKCα maturation by competitively displacing immature PKCα binding to PDK1 during phosphoactivation. Such inhibition of PKCα maturation could slowly lower the cellular concentration of active PKCα, thereby further decreasing PIP_3_ pathway activation by slowing the PKCα-MARCKS-PIP_2_-PI3K-PIP_3_ branch of PIP_3_ production (24, 25). Eventually, however, the falling PKCα levels could remove PKCα inhibition of the PDK1-AKT1 phosphoactivation reaction, providing a feedback control on such PKCα inhibition.

The roles of PKC isoforms in carcinogenesis are complex. In some cases, PKCα may activate the PKCα-MARCKS-PIP_2_-PI3K-PIP_3_ branch of PIP_3_ production (25) and stimulate PIP_3_-PDK1-AKT1-mTOR signaling strongly linked to cell growth and proliferation, thereby providing a potential role for PKC as a cancer/tumor promoter. In other cases, PKC may act as a cancer/tumor suppressor by competitively inhibiting PDK1 activation of AKT1 in the PI3K-PIP3-PDK1-AKT1-mTOR pathway. Evidence for the importance of PKC as a tumor suppressor is provided by reanalysis of the tumor promotion mechanism of phorbol esters including DAG and PMA. Such tumor promoters have recently been shown to deplete PKC populations, indicating that their well-established tumor promotion activity actually stems from loss of PKC as a tumor suppressor (41–43). Due to the variety and complexity of cancer mechanisms, it is plausible that PKC activity may act as either a tumor/cancer promoter or suppressor depending on the cellular context.

More broadly, given the numerous pathways that are downstream of PIP_3_-PDK1-AKT1 signaling, the *cis* mechanism PDK1 phosphoactivation of AKT1 demonstrated by the present findings impacts a wide array of cell processes ranging from cell migration, proliferation, and apoptosis to glucose uptake and glucagon breakdown. Moreover, the findings have general implications for PDK1 regulation of AGC kinases, and for competitive inhibition of AGC kinase signaling by other AGC kinases. Areas for further study include the source of intra-heterodimer flexibility needed to allow an AGC kinase to simultaneously dock to the PDK1 PIF pocket and be phosphoactivated by the PDK1 active site on the opposite face of the PDK1 kinase domain. Such flexibility may plausibly arise from the long, largely unstructured AGC region of the substrate kinase and/or the flexibility of its phosphoactivation loop. Additional studies are needed to delineate the structural basis of the required flexibility, and whether the flexibility is regulated in cells as a novel mechanism for modulation of AGC kinase activation.

## CONCLUSIONS

Overall, the present findings show that membrane-bound PDK1:AKT1 and PDK1:PKCα heterodimers form on a target membrane surface under physiological conditions. Both heterodimers are stabilized by the binding of the AKT1 or PKCα PIF motif, respectively, to the PDK1 PIF pocket, and both are disrupted the small molecule drug PS48 that competitively displaces the PIF motif from the PIF pocket. PKCα exhibits 8-fold tighter PDK1 affinity than AKT1, thus PKCα competitively displaces AKT1 from PDK1:AKT1 heterodimers. Ensemble activity measurements carried out under conditions that closely match those of the single molecule experiments reveal that PDK1 activates AKT1 via a *cis* mechanism by phosphorylating an AKT1 molecule in the same PDK1:AKT1 heterodimer. Moreover, by displacing AKT1 from PDK1, PKCα acts as a strong competitive inhibitor of the PDK1-AKT1 phosphoactivation reaction. Such inhibition suggests that the recently described tumor suppressor activity of PKC (41–43) may arise from its ability to downregulate PDK1-AKT1 phosphoactivation in the PIP3-PDK1-AKT1-mTOR pathway strongly linked to cell growth and oncogenesis.

## AUTHOR CONTRIBUTIONS

Conception JJF, BPZ, MTG; experimental design BPZ, MTG, JJF; data collection MTG, BPZ; data analysis MTG, BPZ; data interpretation JJF, MTG, BPZ; writing of manuscript MTG, JJF; reviewing of manuscript BPZ.

## ACKNOWLEDGEMENTS

The authors gratefully acknowledge funding by NIH Grant R01 GM063235 to JJF, and by NIH Molecular Biophysics Traineeship T32 GM065103 to MTG.

